# Partial derivatives meta-analysis: pooled analyses when individual participant data cannot be shared

**DOI:** 10.1101/038893

**Authors:** Hieab HH Adams, Hadie Adams, Lenore J Launer, Sudha Seshadri, Reinhold Schmidt, Joshua C Bis, Stephanie Debette, Paul A Nyquist, Jeroen Van der Grond, Thomas H Mosley, Jingyun Yang, Alexander Teumer, Saima Hilal, Gennady V Roshchupkin, Joanna M Wardlaw, Claudia L Satizabal, Edith Hofer, Ganesh Chauhan, Albert Smith, Lisa R Yanek, Sven J Van der Lee, Stella Trompet, Vincent Chouraki, Konstantinos A Arfanakis, James T Becker, Wiro J Niessen, Anton JM de Craen, Fabrice F Crivello, Li An Lin, Debra A Fleischman, Tien Yin Wong, Oscar H Franco, Katharina Wittfeld, J Wouter Jukema, Philip L De Jager, Albert Hofman, Charles DeCarli, Dimitris Rizopoulos, WT Longstreth, Bernard M Mazoyer, Vilmundar Gudnason, David A Bennett, Ian J Deary, M Kamran Ikram, Hans J Grabe, Myriam Fornage, Cornelia M Van Duijn, Meike W Vernooij, M Arfan Ikram, on behalf of the HD-READY Consortium

**Affiliations:** Department of Epidemiology, Erasmus MC, the Netherlands.; Department of Radiology, Erasmus MC, the Netherlands.; Division of Neurosurgery, Department of Clinical Neurosciences, Addenbrooke’s Hospital & University of Cambridge, UK; Intramural Research Program, National Institute on Aging, USA; Department of Neurology, Boston University School of Medicine, USA; Department of Neurology, Medical University Graz, Austria; Cardiovascular Health Research Unit, Department of Medicine, University of Washington, USA; INSERM U897, University of Bordeaux, Bordeaux, France; University of Bordeaux, Bordeaux, France; Department of Neurology, Boston University School of Medicine, Boston, MA, USA; Department of Neurology, Bordeaux University Hospital, Bordeaux, France; Department of Neurology, Johns Hopkins, USA; Department of Anesthesia/Critical Care Medicine, Johns Hopkins, USA; Department of Neurosurgery, Johns Hopkins, USA; Department of Radiology, Leiden University Medical Center, Netherlands; Department of Medicine, University of Mississippi Medical Center, USA; Rush Alzheimer’s Disease Center, Rush University Medical Center, USA; Department of Neurological Sciences, Rush University Medical Center, USA; Institute for Community Medicine, University Medicine Greifswald, Germany; Department of Ophthalmology, National University of Singapore, Singapore; Department of Pharmacology, National University of Singapore, Singapore; Department of Medical Informatics, Erasmus MC, the Netherlands; Centre for Clinical Brain Sciences, University of Edinburgh, UK; Centre for Cognitive Ageing and Cognitive Epidemiology, University of Edinburgh, UK; Institute for Medical Informatics, Statistics and Documentation, Medical University Graz, Austria; Icelandic Heart Association, Iceland; Department of General Internal Medicine, Johns Hopkins, USA; Department of Cardiology, Leiden University Medical Center, Netherlands; Department of Gerontology and Geriatrics, Leiden University Medical Center, Netherlands; Department of Biomedical Engineering, Illinois Institute of Technology, USA; Diagnostic Radiology and Nuclear Medicine, Rush University Medical Center, USA; Department of Psychiatry, Neurology and Psychology, University of Pittsburgh, USA; CNRS UMR5296, University of Bordeaux, France; Human Genetics Center and Institute of Molecular Medicine, University of Texas Health Science Center at Houston, USA; Duke-NUS Graduate Medical School, National University of Singapore, Singapore; Singapore Eye Research Institute, Singapore National Eye Centre, Singapore; German Center for Neurodegenerative Diseases (DZNE), Site Rostock/Greifswald, Germany; Durrer Center for Cardiogenetic Research, Amsterdam, The Netherlands; Department of Neurology & Psychiatry, Brigham and Women’s Hospital, USA; Program in Medical and Population Genetics, Broad Institute, Cambridge, MA, USA; Harvard Medical School, USA; Department of Neurology, UC Davis, USA; Center for Neuroscience, UC Davis, USA; Department of Biostatistics, Erasmus MC, The Netherlands; Department of Neurology, University of Washington, USA; Department of Epidemiology, University of Washington, USA; Faculty of Medicine, University of Iceland, Iceland; Department of Psychology, University of Edinburgh, UK; Memory Aging & Cognition Centre, National University of Singapore, Singapore; Department of Psychiatry & Psychotherapy, University Medicine Greifswald, Germany; Department of Neurology, Erasmus MC, the Netherlands

## Abstract

Joint analysis of data from multiple studies in collaborative efforts strengthens scientific evidence, with the gold standard approach being the pooling of individual participant data (IPD). However, sharing IPD often has legal, ethical, and logistic constraints for sensitive or high-dimensional data, such as in clinical trials, observational studies, and large-scale omics studies. Therefore, meta-analysis of study-level effect estimates is routinely done, but this compromises on statistical power, accuracy, and flexibility. Here we propose a novel meta-analytical approach, named partial derivatives meta-analysis, that is mathematically equivalent to using IPD, yet only requires the sharing of aggregate data. It not only yields identical results as pooled IPD analyses, but also allows post-hoc adjustments for covariates and stratification without the need for site-specific re-analysis. Thus, in case that IPD cannot be shared, partial derivatives meta-analysis still produces gold standard results, which can be used to better inform guidelines and policies on clinical practice.

## Introduction

Science is an increasingly collaborative effort. The benefits of jointly investigating research questions, such as improved power to detect effects and generalizability of results,^1-3^ have been known for long, but the recent technology-driven emergence of high-dimensional datasets, e.g. in the fields of genetics and imaging, has further underlined the need for collaboration.^4^ The limited capacity of single studies to collect these data in sufficient numbers has resulted in the formation of numerous large consortia.^5-7^

Generally, joint analyses are performed by either pooling individual participant data (IPD) from multiple studies or by meta-analysing aggregate data that are derived from study-specific analyses. In pooled analyses, raw participant data are shared to produce combined datasets that are then analysed as a single study. In conventional aggregate data meta-analysis, only the study-specific effect estimates are provided or extracted from literature and these estimates are averaged to approximate the overall effect for all the studies combined. Although pooled analyses provide the highest power and accuracy,^4, 8, 9^ several barriers exist to sharing IPD, including when local legislation prevents the release of collected data, when the process of sharing data presents logistic problems due to its size, or when the data, despite being anonymized, can lead to the identification of individuals and reveal sensitive medical information.^10^ Even though the research community as well as funding agencies put considerable pressure on investigators to make datasets publicly available, in these conditions, IPD cannot be pooled for joint analysis.

Here we propose a novel meta-analytical approach, *partial derivatives meta-analysis*, which 1) provides the statistical and analytical benefits of pooled analyses for linear regression models, 2) uses aggregate data that cannot be traced back to individual participant data, and 3) is easily applicable in current research settings. We use examples from clinical trials, observational studies, and high-dimensional omics to illustrate the broad relevance of this approach.

## Methods

### Regression analysis

Regression models estimate the effects of predictor variables on the outcome variable by determining the combination of effect estimates that correspond best to the data at hand. When performing a regression analysis, the IPD is converted into several intermediate statistics before arriving at the eventual results, i.e. the effect estimates (**Figure 1**). First, residual error terms are calculated for each individual. This error represents the difference between the actual outcome variable and its value predicted by the model. Second, the error terms are consolidated into a set of partial derivatives equations, which are the sum of all individual terms. Third, the partial derivatives equations are set to zero and solved, to find the best combination of effect estimates that minimize the error, i.e. give a predicted value as close to the actual outcome variable. The “fitting of the model” refers to this final step.

**Figure 1:**
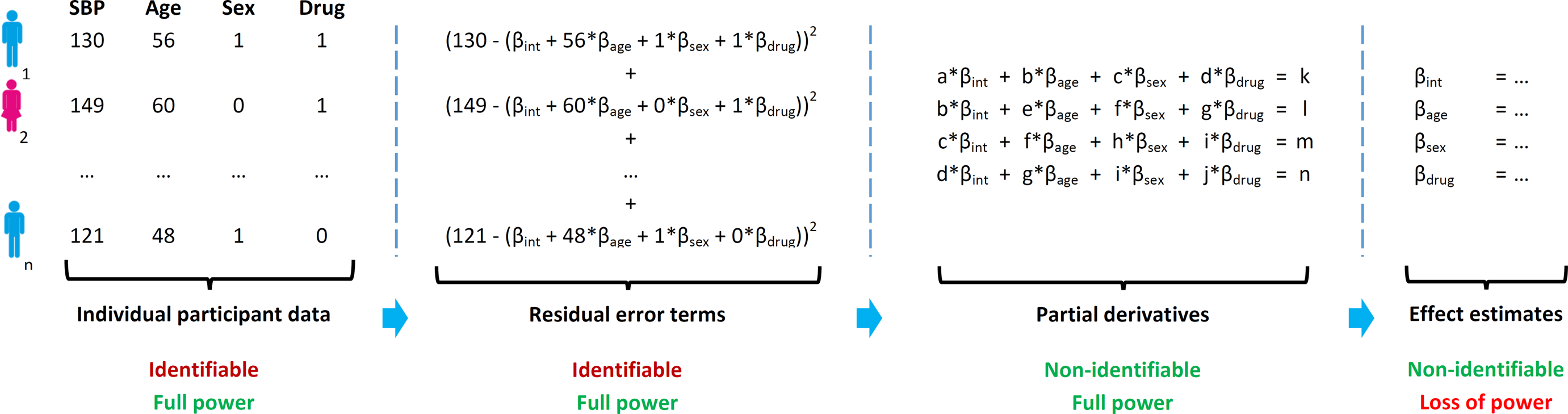
Four stages of data in linear regression analysis and their meta-analytical properties when shared. Schematic overview of four stages when performing a linear regression analysis for a study with *n* participants, where systolic blood pressure (SBP) is modelled using predictors age, sex, and a novel drug of interest. Starting with the individual participant data, the residual error terms are calculated for each individual separately. Subsequently, the partial derivatives are generated by adding up the residual error terms of all individuals. Finally, simultaneously solving the partial derivatives gives the effect estimates for each of the predictors and the intercept per study. When sharing the individual participant data, residual error terms, or partial derivatives, it is possible to calculate meta-analysed effect estimates without the loss of power. Only by exchanging the partial derivatives or effect estimates, however, is the shared information not identifiable to a participant level.

### Partial derivatives meta-analysis

Although the partial derivatives are only intermediate statistics for calculating effect estimates, they have great potential for joint analyses, in particular with linear regression. In linear regression, partial derivatives are calculated by multiplying every variable with all the others, resulting in one value for every pair of variables. This is done separately for each participant in the analysis and these participant-specific values are subsequently summed together to obtain the partial derivatives (**Figure 2A**). The process of summation can be split for different groups of participants and combined afterwards, without compromising the end result. Extending from this concept, partial derivatives could be calculated separately for a group of participants (i.e., within different studies) and summed together at the meta-analytical stage—a strategy we term *partial derivatives meta-analysis* (**Figure 2B;** details in the **Supplementary Material**). Using these summed partial derivatives it is then possible to fit a (meta-analysed) model, which yields effect estimates that are mathematically identical to those from pooled analyses, without the sharing of IPD. Additionally, this approach allows for post-hoc changes in the covariate adjustments (e.g., removing the variable “age” from the model) and stratified analyses (e.g., studying men and women separately) without site-specific re-analyses.

**Figure 2:**
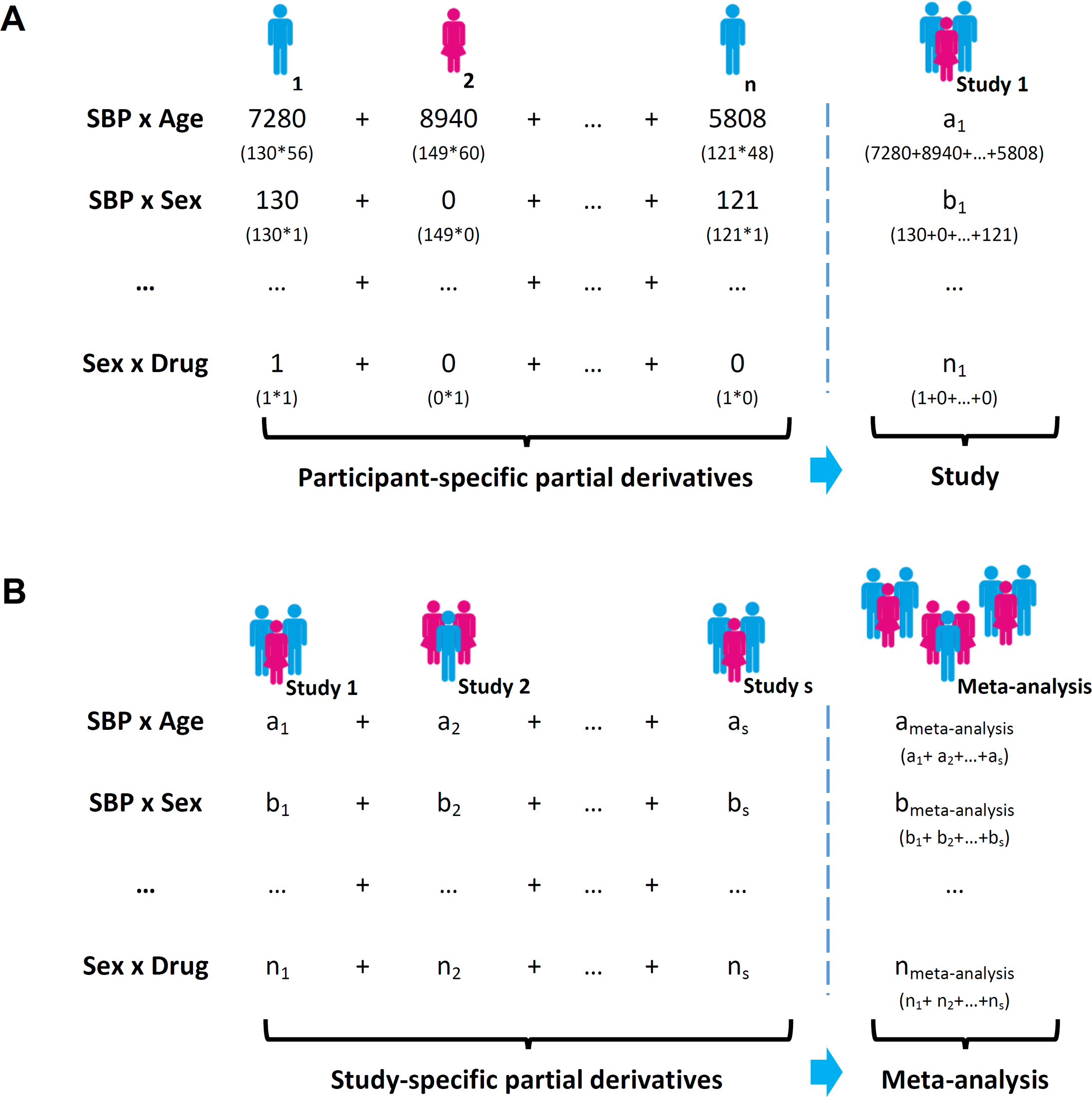
Strategy for performing a partial derivatives meta-analysis. Overview of calculation for a linear regression analysis for a meta-analysis of s studies, where systolic blood pressure (SBP) is modelled using predictors age, sex, and a novel drug of interest. For each of *n* participants in the first study, partial derivatives are calculated by multiplying the value of every variable with the value of all the others, resulting in a single value for every pair of variables (panel A, left side). The participant-specific values are subsequently summed together to obtain the partial derivatives of the whole first study (panel A, right side). Analogous to this, the partial derivatives can be calculated for each study separately (panel B, left side) and combined at the meta-analytical stage (panel B, right side). The resulting partial derivatives are mathematically equivalent to those that would have been obtained if all individual participant data was available and therefore provide identical results.

## Results

In this section, we show how partial derivatives meta-analysis can be applied in current research settings and compare this to the conventional methods, namely the sharing of IPD (gold standard) and the sharing of effect estimates. To illustrate the broad applicability of this approach, we use examples from clinical trials, observational studies, and omics studies.

### Descriptive statistics

Papers describing investigations in humans generally start with a table containing the (baseline) characteristics of the study population. Using IPD, it is easy to generate these descriptive statistics, but this is not possible using the aggregated effect estimates. Instead, all contributing sites need to calculate their descriptive statistics locally, leading to a potential source of errors (e.g., using different individuals than for the main analysis).

The partial derivatives are a set of values, the exact number of which depends on the number of predictor variables in the chosen model (see **Supplementary Material**), and would for example equal 14 values when modelling 4 predictors. Interestingly, some of these values can be used to determine descriptive statistics, such as the sample size, but also the means and standard deviations of any of the variables, including the outcome. This therefore obviates the need for sites to provide these descriptive statistics separately.

### Full statistical power: example using omics data

Statistical power to detect associations is important for all studies, but its relevance is perhaps most apparent for large-scale omics studies such as genome-wide association studies. Here, strict multiple testing correction for millions of tests and the small effect sizes of genetic variants make it difficult to identify true associations, and this has necessitated collaborative efforts. Pooling IPD is the gold standard and yields the greatest power, although conventional meta-analysis of effect estimates can provide similar results in specific cases.^11, 12^ However, when some sites cannot share their IPD, this leads to a smaller sample size than would be obtained using conventional meta-analysis, and this is even more detrimental to the statistical power. In comparison, partial derivatives meta-analysis has the potential to overcome both limitations.

First, using simulated data, we show that the effect estimates of partial derivatives meta-analysis are identical to those from a pooled IPD analysis, while this is not the case for conventional meta-analysis (**Figure 3A**). Additionally, partial derivatives meta-analysis increasingly outperformed conventional meta-analysis when, for a fixed total sample size, the number of studies were increased (i.e., the size of individual studies was smaller) (**Figure 3B**). For conventional meta-analysis, small sample sizes or complex models results in unstable (and eventually unobtainable) effect estimates within each site, which generally leads to the exclusion of these sites. Besides the statistical loss of power, this selection forms an epidemiological problem in the light of the biases associated with leaving out particular studies. In contrast, partial derivatives meta-analysis does not require a minimum number of participants and could theoretically even be performed on a single individual, although in this case the participant data would of course be identifiable.

**Figure 3:**
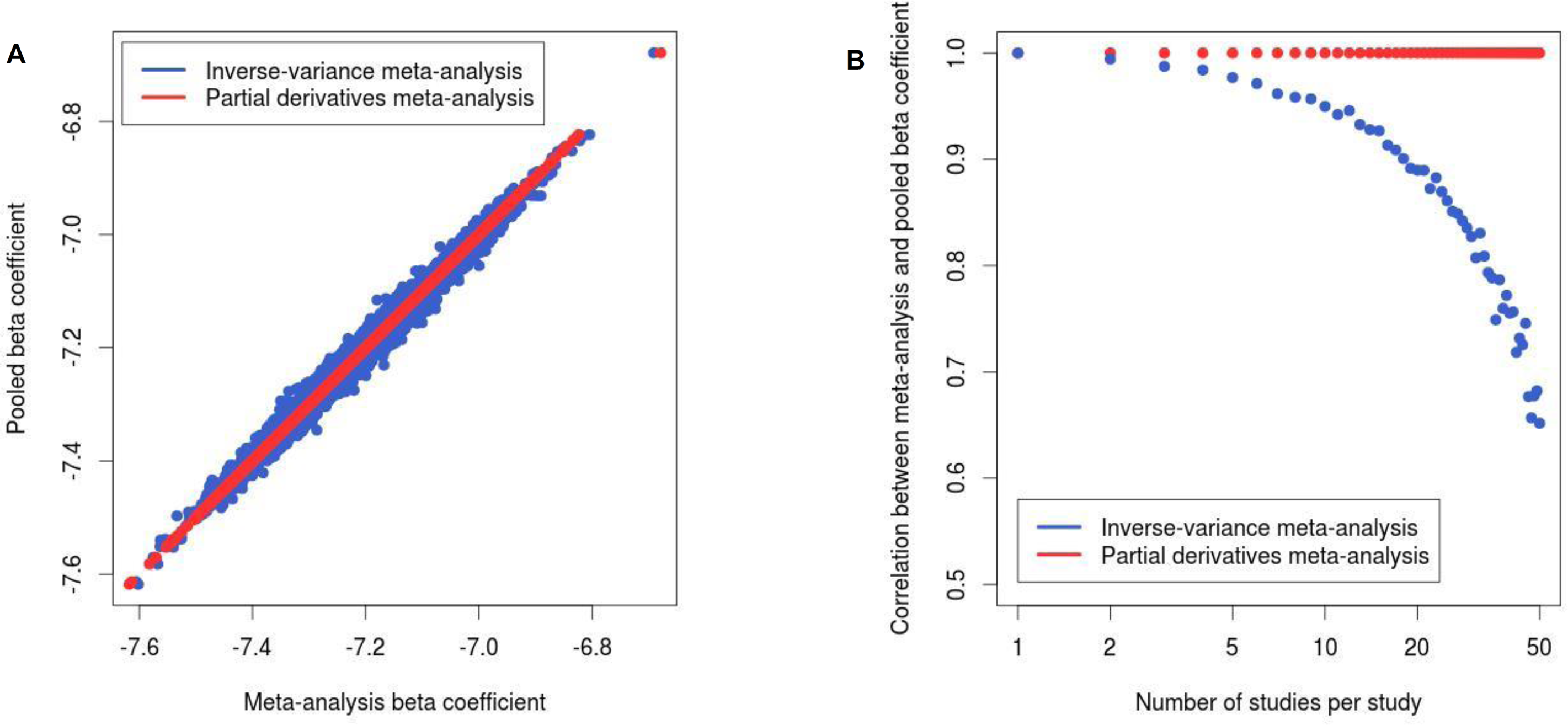
Correlation of effect estimates from conventional and partial derivatives meta-analysis with effect estimates from a pooled analysis. Panel A show a scatter plot of beta coefficients obtained from inverse-variance meta-analysis of effect estimates and pooled analysis (royal blue) and from partial derivatives meta-analysis and pooled analysis (firebrick red). Results were obtained from simulation with the following parameters: number of studies = 2, total sample size = 500, beta coefficient of predictor = −7.20, number of simulations = 10,000. Panel B shows correlation coefficients of beta coefficients between inverse-variance meta-analysis of effect estimates and a pooled analysis (royal blue) and between partial derivatives meta-analysis and a pooled analysis (firebrick red), with a varying number of studies and study size. Simulation parameters were as follows: total sample size = 500, number of studies = 1 to 50, number of simulations = 10,000. All simulations were performed in R (version 3.0.1)

Second, using real data of 14,643 participants from 19 cohort studies from the HD-READY consortium (see **Supplementary Material**), we show that the anonymised nature of partial derivatives can overcome the data sharing obstacles related to IPD. Of the 19 participating studies, only 4 were able to share IPD with investigators from the meta-analytical site, resulting in a pooled IPD analysis of 5,611 individuals, less than 40% of the total sample (**Table 1**). In contrast, all studies were able to contribute partial derivatives. Studies sharing IPD did not need to perform any on-site analyses. For studies running partial derivatives (which were all studies), all site-specific data could be generated through a single analysis. Studies sharing effect estimates needed to perform additional analyses for every model, thus increasing the analytical burden.

**Table 1.** Comparison of sharing conventional data and partial derivatives with respect to study site participation and analytical possibilities for a proof-of-principle study.

| Characteristic | IPD | Partial derivatives | Effect estimates |
| --- | --- | --- | --- |
| **Study characteristics** |  |  |  |
| Number of studies agreed to participate | 7<sup>§</sup> | 19 | 19 |
| Number of participants | 5,611 | 14,643 | 14,643 |
| Country of origin |  |  |  |
| Netherlands | 5 | 5 | 5 |
| United Kingdom | 0 | 1 | 1 |
| Germany | 0 | 2 | 2 |
| France | 0 | 1 | 1 |
| Austria | 1 | 1 | 1 |
| Iceland | 0 | 1 | 1 |
| Singapore | 1 | 1 | 1 |
| United States | 0 | 7 | 7 |
| **Data sharing and analysis** |  |  |  |
| Sharing of non-identifiable data | No | Yes | Yes |
| Individual-level effect estimates | Yes | Yes | No |
| Stratified analyses | Yes | Yes | No |
| Post-hoc model adjustments | Yes | Yes | No |
| Number of required analyses per site<sup>†</sup> | 0 | 1 | 21 |

† Calculated as number of analyses required for seven different models, for the complete sample as well stratified for sex.
§ These are the meta-analysis sites (Rotterdam Study 1-3), ASPS-Fam, EDIS, LLS, and PROSPER.

Third, using the data from these 19 studies, we compared the different meta-analytical methods to investigate the association between an established genetic risk variant (rs9939609, *FTO/IRX3* locus) and body mass index (**Figure 4**). Conventional meta-analysis of effect estimates showed a genome-wide significant association (ß = 0.34, P = 2.19 × 10^−11^). Next, we confirmed that partial derivatives meta-analysis indeed yields identical results as pooled analyses in the subset of participating cohorts that were able to share their IPD. We then used the partial derivatives to calculate the effect estimates under various scenarios of pooled analyses. Naïve pooling of samples, which assumes a completely homogeneous population, resulted in biased estimates (ß = 0.47, P = 4.94 × 10^−18^), whereas controlling for study site using an intercept for each cohort resolved this issue (ß = 0.34, P = 4.95 × 10^−11^). Modelling cohort-specific effects for all the covariates in the model (intercept, age, and sex) provided a slightly larger effect estimate that was more significant than conventional meta-analysis (ß = 0.35, P = 5.17 × 10^−12^). At first sight, this improvement over the conventional meta-analysis is modest. However, large-scale omics studies routinely involve millions of analyses, and the increase in power for such investigations will thus be cumulative.

**Figure 4:**
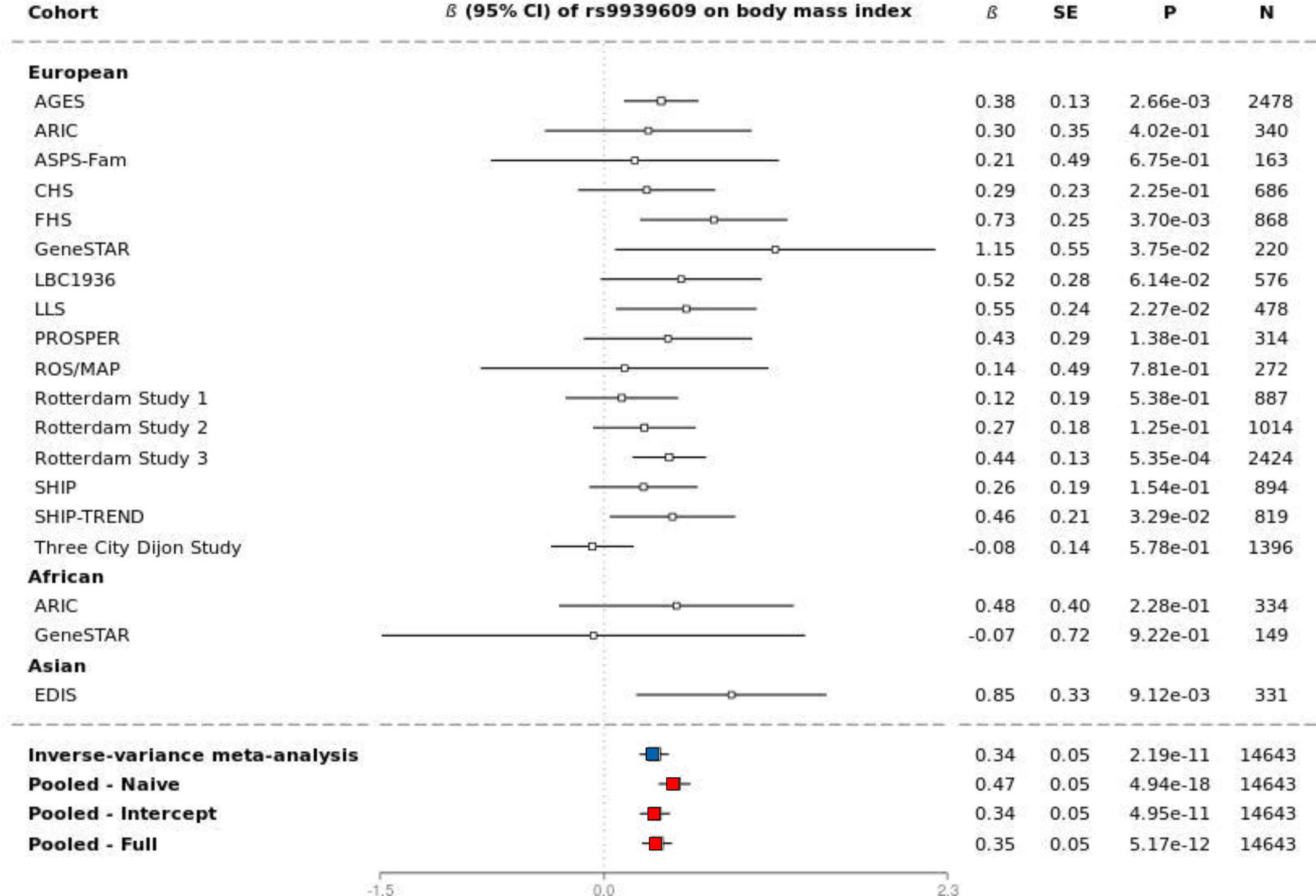
Forest plot for the association of rs9939609 with body mass index. Forest plots showing beta coefficients and 95% confidence intervals of individual cohorts as well as four different joint analyses, all obtained from the partial derivatives: Inverse-variance based meta-analysis of effect estimates, pooling without additional adjustments (“Naïve”), pooling with adjustment for cohort (“Intercept”), and pooling with adjustment for cohort and other cohort-specific covariates (“Full”).

### Post-hoc adjustment of the model: example using observational data

Although the increase in power of partial derivatives meta-analysis is advantageous, the increased flexibility of the method is particularly appealing. Similar to an IPD analysis, partial derivatives meta-analysis allows for adjustments of statistical models without the need for each contributing study to reanalyse its data and share new results. All predictors of interest, including those to be added or removed in a later stage, can simply be included when generating the partial derivatives; when fitting the model on the meta-analytical level, the decision can be made about what specific predictors to include. For conventional meta-analysis, post-hoc adjustments are not possible, as this would require re-analyses from each site. We illustrate this application using data from three observational cohorts (total N = 5,431), where the aim of the study is to understand how body height relates to general cognitive function.

In this study, we defined general cognitive function as the common variance between multiple cognitive tests, as previously described.^13^ The pooled age- and sex-adjusted analysis showed a strong association between body height and cognitive function (ß = 0.016, P = 3.03 × 10^−15^). However, given that body height is an indicator of educational attainment and this in turn is related to cognitive function, it remains unclear whether this association is independent of education. With the partial derivatives, education can be added to the model at the meta-analytical level to test this and thereby provide more information for correct inference of the results. Adjustment for education resulted in an attenuated, but still significant, effect estimate of height (ß = 0.010, P = 9.78 × 10^−8^).

Naively pooling IPD from all cohorts together does not sufficiently account for any underlying heterogeneity that may exist between the cohorts.^14, 15^ For example, the education variable was not on the same scale for all three cohorts due to the use of a different questionnaire, and it might not be appropriate to treat them the same. Conventional meta-analysis, on the other hand, assumes complete heterogeneity among the cohorts for all variables in the model. However, using partial derivatives, these assumptions can be formally tested and modelled on the meta-analytical stage. It is possible to determine the study-specific effects for all variables, and also determine whether some studies cluster together. In our example, we found that the effects of education were indeed significantly different between the cohorts, whereas the intercept did not differ between two of the cohorts, which were demographically similar.

### Stratified analyses: example using clinical trial data

Another application of partial derivatives is the possibility to perform stratified analyses. For a comparative clinical trial, stratification in the form of subgroup analyses provides means of investigating whether certain patient groups are likely to respond differently to a treatment. Consider randomized, clinical trials that aim to determine the efficacy of a novel blood pressure lowering medication. If the treatment is found to be successful, it might be interesting to see if the benefit is the same for men and women. In case of a negative trial, it could be informative to find that persons with kidney disease do not tolerate the treatment, whereas there is a beneficial effects in those without. If the subgroups are pre-specified, the partial derivatives will contain the necessary information to perform these joint analyses of the clinical trials without exchanging the IPD.

## Discussion

We show that sharing of partial derivatives among studies is sufficient to calculate meta-analysed effect estimates for linear regression models that are mathematically equivalent to a pooled analysis, the gold standard, without the need to share IPD. Partial derivatives meta-analysis outperforms conventional meta-analysis, in particular for complex models and with an increasing number of (smaller) studies. Furthermore, partial derivative meta-analysis has additional advantages over conventional meta-analysis, including the ability to perform post-hoc adjustments of the model and detailed subgroup analyses, as well as the possibility to include small studies in a meta-analysis that would otherwise have been excluded. For the specific case of high-dimensional data, sharing of partial derivatives is in fact more efficient than sharing IPD given the size of the data.

Thus, the partial derivatives are in fact ‘pluripotent’ since not only IPD effect estimates can be calculated, but also those from conventional meta-analysis. Additionally, this approach enables the examination of each of the predictors separately and for different combinations of studies that might cluster together. Therefore, in cases where naïve pooling of IPD might not be appropriate, a different analytic approach will be possible using the same partial derivatives. The increased flexibility of sharing partial derivatives also requires increased caution. Just as with sharing of IPD and effect estimates, the partial derivatives can be used for analyses other than those of the primary research question. Subgroup analyses in clinical trials require rigorous reporting,^16, 17^ and we suggest to follow these standards to prevent secondary hypothesis testing not considered in the primary analysis plan.

A main focus when developing this method was its ease of use for other researchers. To promote the dissemination of partial derivatives meta-analysis, we provide examples for calculating the partial derivatives with commonly used statistical software (R, SPSS, Excel, SAS, and STATA). Additionally, we have implemented partial derivatives meta-analysis in HASE, a software package for genome-wide association studies.^18^

Here we described a meta-analytical approach for continuous outcomes using linear regression analysis, but additional work is needed for applying this approach to binary outcomes and time-to-event data. The usual method for parameter estimation in generalized linear models and proportional hazards models is through multiple rounds of individual-level calculations to find the maximum likelihood numerically; a non-iterative approach would therefore be more feasible to reduce demands on contributing studies. Other proposed solutions for pooling of individual-level data include projects such as BioSHaRE and DataSHIELD.^17, 19, 20^ In these projects, for each analysis, summary statistics are continuously exchanged between servers of different sites until an adequate model is fit. Although this is promising from a methodological perspective, it does require rigorous data harmonization and subsequent linking of the data to external servers using dedicated software packages. The high level of collaboration needed between the studies and the time-investment for preparation of the data could potentially explain the limited implementation by other researchers.

Also, using DataSHIELD or conventional aggregate data, published results cannot be reused for calculating effect estimates identical to IPD meta-analysis. With partial derivatives meta-analysis however, the partial derivatives can be provided and, with new studies, added to get new estimates. We therefore propose publishing of partial derivatives in addition to the conventional effect estimates, while retaining IPD within host institutions of participating studies when this cannot be made available too.

Interestingly, many of the partial derivatives values correspond to descriptive statistics already being published on a frequent basis (e.g. sample size, mean age, number of women), along with the effect estimates for the variables of interest. This raises the intriguing possibility of using published results to recalculate the partial derivatives and subsequently perform a pooled analysis. Not all values of the partial derivatives are available, however, so some assumptions would need to be made about the correlation between variables (e.g., by using data from an available sample). This possibility is especially interesting for published data for which it is not feasible to request re-analysis, e.g. when investigators are not reachable or they do not wish to participate.

Another potentially useful avenue is for computationally intensive analyses that are common in the omics field, such as genome-wide association studies. Partial derivatives meta-analysis only requires a “partial” regression to be performed within each site, since the partial derivatives are needed for sharing, but it is not necessary to subsequently calculate the effect estimates. The fitting of the model will be done only once, after combining all the partial derivatives from the contributing sites, thereby shifting part of the computational burden to the stage of the meta-analysis. Also, it is possible to use the same partial derivatives to perform exploratory analyses, for instance additional adjustments for potential confounders, which are particularly time-consuming for large-scale omics studies, and therefore are often not performed. However, since these exploratory analyses can be instrumental for the correct causal inference of findings, the use of partial derivatives meta-analysis could facilitate the completion of scientific efforts and potentially accelerate scientific discoveries. Furthermore, in addition to the usual difficulties accompanying the exchange of IPD, the sharing of aggregate data actually outperforms the sharing of IPD with respect to the time needed for transferring the files,^21^ making this approach more efficient and perhaps less costly.

Ultimately, sharing IPD is still superior with respect to flexibility and insight into data. However, until such sharing is adopted by the scientific community as a whole, partial derivatives meta-analysis could provide a way forward while using only aggregate data.

## Authors’ contributions

HHHA conceived of the study, participated in its design, performed the statistical analysis, interpreted the data, and drafted the manuscript. HaA conceived of the study, interpreted the data, and helped draft the manuscript. JCB, PAN, JvdG, JY, AT, SH, CLS, EH, GC, AS, LRY, SJvdL, ST, VC, FFC, LAL, and TYW performed statistical analysis and revised the manuscript critically for important intellectual content. LJL, SS, RS, SD, THM, JMW, KAA, JTB, WJN, AJMdC, DAF, KW, JWJ, PLDJ, AH, CD, WTL, BMM, WG, DAB, IJD, MKI, HJG, MF, and CMvD acquired data and revised the manuscript critically for important intellectual content. GR, OHF, and DR, interpreted the data and revised the manuscript critically for important intellectual content. MWV and MAI participated in its design, interpreted the data, and helped draft the manuscript. All authors read and approved the final manuscript.

## Authors’ information

The authors of this manuscript participate in multiple large-scale collaborations such as the CHARGE, ENIGMA, and UNIVRSE consortia. Their experiences with data sharing, particularly the encountered obstacles that hinder the best possible research, have led to the investigation of alternative approaches of joint analyses and eventually to the development of the current method, partial derivatives meta-analysis.

## Acknowledgements

**HD-READY consortium:** This project is funded by the EU Joint Programme - Neurodegenerative Disease Research (JPND).

The cohort-specific acknowledgements can be found in the **Supplementary Material**.

